# Cardiac disease diagnosis based on GAN in case of missing data

**DOI:** 10.1101/2023.09.22.559043

**Authors:** Xing Chen, Na Zhang, Xiaohui Yang, Chunyan Wang, Na Qi, Tianyun Luan, Wendi Zhu, Chenjie Zhang, Chao Yang

## Abstract

In daily life, two common algorithms are used for collecting medical disease data: data integration of medical institutions and questionnaires. However, these statistical methods require collecting data from the entire research area, which consumes a significant amount of manpower and material resources. Additionally, data integration is difficult and poses privacy protection challenges, resulting in a large number of missing data in the dataset. The presence of incomplete data significantly reduces the quality of the published data, hindering the timely analysis of data and the generation of reliable knowledge by epidemiologists, public health authorities, and researchers.

Consequently, this affects the downstream tasks that rely on this data. To address the issue of discrete missing data in cardiac disease, this paper proposes the AGAN (Attribute Generative Adversarial Nets) architecture for missing data filling, based on generative adversarial networks. This algorithm takes advantage of the strong learning ability of generative adversarial networks. Given the ambiguous meaning of filling data in other network structures, the attribute matrix is designed to directly convert it into the corresponding data type, making the actual meaning of the filling data more evident. Furthermore, the distribution deviation between the generated data and the real data is integrated into the loss function of the generative adversarial networks, improving their training stability and ensuring consistency between the generated data and the real data distribution. This approach establishes the missing data filling mechanism based on the generative adversarial networks, which ensures the rationality of the data distribution while filling the missing data samples. The experimental results demonstrate that compared to other filling algorithms, the data matrix filled by the proposed algorithm in this paper has more evident practical significance, fewer errors, and higher accuracy in downstream classification prediction.

## Introduction

### Background

Cardiac disease is a major global health problem that affects millions of people every year. With the increasing prevalence of cardiac disease, early detection and accurate prediction of its occurrence are crucial for effective management and treatment.

Advances in computer science and technology have provided new opportunities for the medical field to address this problem. Machine learning and artificial intelligence algorithms have been applied to analyze large datasets of medical information to develop predictive models for cardiac disease [1]. According to data from the United States Centers for Disease Control and Prevention (CDC), cardiac disease remains a leading cause of death for most races in the United States, including African Americans, American Indians, Alaska Native Americans, and whites [2].

Approximately 47% of Americans have at least three of the key risk factors for cardiac disease, such as high blood pressure, high cholesterol, and smoking, among others.

Other important indicators include diabetes status, obesity (measured by high BMI), physical inactivity, and excessive drinking [3]. In China, the Interpretation of Key Points of China’s Cardiovascular Health and Disease Report 2021, released in 2021, clearly points out that the number of people suffering from cardiovascular diseases is on the rise, with rates of 45.91% in rural areas and 43.56% in urban areas. Early diagnosis and timely treatment are essential measures to address this crisis. Moreover, the incidence of cardiac disease in cardiovascular disease centers is increasing and gradually showing a younger trend, which has raised health alarms for more young people [4].The economic burden of cardiovascular diseases on residents and society is also increasing, making it a major public health problem. Therefore, it is crucial to strengthen government-led prevention and treatment of cardiovascular diseases [5].

Cardiovascular disease is associated with high disability and mortality rates. Despite the advanced treatment methods available at present, many patients remain unable to care for themselves after being cured. As such, primary prevention of cardiovascular disease is particularly important. Primary prevention involves modifying unhealthy living habits before the onset of the disease, such as quitting smoking, adopting a balanced diet, increasing physical activity, and taking medications to control risk factors such as blood pressure, cholesterol levels, and blood sugar. Although the pathogenesis of cardiovascular diseases has not been fully determined, a significant number of studies have confirmed that all types of cardiovascular diseases share a common pathological basis, and the primary risk factors have been identified. Most of these factors can be managed through lifestyle modifications. Therefore, it is crucial to develop accurate and efficient early prediction tools for cardiovascular diseases to identify high-risk groups and provide early warnings. Furthermore, it is recommended that high-risk individuals change their unhealthy living habits to control risk factors and reduce the risk of developing cardiovascular disease [6]. However, at this stage, many rare diseases have limited data samples. Typically, people only seek medical attention when they experience discomfort. Take cancer, for example; by the time people experience discomfort, the tumor has often formed and may have already progressed to the middle or late stage of the disease. Before targeted drugs corresponding to cancer become available, doctors can only examine the patient’s DNA data and cluster them based on the detected data, assuming that the number of clusters is known in advance. Once the clustering results are obtained, different treatment methods are implemented based on the clustering results, as medical resources are valuable. However, the cost of DNA testing is high, and the number of patients with certain types of cancer is relatively small, making diagnosis more challenging.

### Related work

Traditional algorithms for processing missing data are mainly divided into two categories: deletion and filling. The deletion method involves obtaining a “complete” data set by directly deleting missing items. Its advantage is that it is simple and easy to operate; however, this method leads to information loss and deviation in data distribution, rendering it only applicable to cases where there is a small proportion of missing items. The filling method, on the other hand, can be divided into statistics-based filling and model-based filling. The former includes mean filling, random filling, forward (backward) interpolation, weighted filling, and other methods that utilize the statistical characteristics of the missing data to fill in the gaps. These algorithms yield stable filling results and are applicable to situations where the distribution characteristics of the missing sequence are simple and clear, and the variable correlation is strong. The model-based filling methods, such as regression model filling, K-nearest neighbor interpolation, and the expectation maximization algorithm (EM) filling, involve selecting different models or algorithms to measure the distance between missing and observed data. The goal is to adjust the parameters to minimize this deviation and estimate missing data. This approach makes full use of the information contained in the existing data, but has specific requirements for the data set itself, such as strong correlations and a specific distribution of the data.

Overall, there are various algorithms for processing missing data, each with its own advantages and limitations. The selection of an appropriate method depends on the characteristics of the data set and the specific research question.

Numerous algorithms have been proposed for filling missing data, with discriminant and generative algorithms being the two advanced approaches. Discriminant algorithms include chain equation multiple filling algorithm, random forest algorithm, and matrix filling algorithm, while generative algorithms include expectation maximization algorithms and deep learning-based algorithms, such as noise reduction automatic encoders (DAE) [7] and generating adversarial networks (GAN) [8]. However, some of these algorithms have limitations. For instance, Fedus [9] et al. generate timing data based on self-coder, but the training process of this algorithm requires complete timing data, which makes it difficult to guarantee the generation effect when the training data is missing. Gheyas [10] et al. solved the filling problem through denoising self-coder, but their algorithm uses the average value or the most commonly used label to replace the missing data when initializing the network, which may distort the multivariate relationship. Che [11] et al. developed the GRU-D model, which achieves better prediction results, but cannot be used for unsupervised learning without prediction tags. Yoon [12] et al. applied the generation countermeasure network to the generation of missing data but did not take into account the dependency between the time series data. Zheng [13] et al. proposed a fault prediction method based on double GAN, which is mainly aimed at the situation where the training samples are small but the data set is continuous and complete.

Luo [14] et al. used GRU to form GAN to realize the generation of time series data, but this algorithm cannot guarantee that the filling result of missing data conforms to the overall distribution. Therefore, while various algorithms have been developed for filling missing data, each algorithm has its limitations and is best suited to certain types of data sets.

The EM algorithm for missing data imputation was first proposed in 1977 by Dempster, Laird, and Rubin, and has since attracted significant attention and research from scholars [15]. It is an iterative algorithm based on maximum likelihood estimation, which simplifies a complex estimation problem by assuming the existence of hidden variables and simplifying the likelihood equation. Through subsequent improvements and applications, such as the ECM algorithm [16] proposed by Meng and Rubin, and the MCEM algorithm [17] that combines Monte Carlo simulation with the E-step, the EM algorithm has become a widely used and stable parameter estimation method for missing data imputation. Recent research has expanded the methods for missing data imputation from simple methods like mean filling and Hotdecking to more complex model-based methods like regression filling, statistical learning methods such as missing forest filling and automatic encoding filling. These advancements have reduced the requirements for data hypothesis and improved the accuracy of missing data imputation.

It is true that there are not many technical means to fill the missing data based on deep learning, and the research on filling the missing data using Generative Adversarial Networks is still relatively new. However, GAN has shown great potential in generating realistic data and filling missing data [18]. GAN has been used for image generation and image reconstruction, and these tasks are similar to missing data filling, as they both involve learning the underlying distribution of data and generating data sets subject to sample distribution under certain constraints. Unlike traditional missing data filling methods that rely on strict assumptions, GAN does not have any explicit assumptions about data, and they can learn complex relationships and patterns in the data. The larger the amount of data, the more accurate the results will be, as GAN can use the additional data to improve the accuracy of the generated data. However, GAN can also face challenges such as mode collapse or gradient disappearance during the training process [19], which can affect the quality of the generated data.

### Our work

The algorithms mentioned above have yielded promising results in the context of data filling. However, certain issues still need to be addressed. Firstly, the error between the filled data and the real data is substantial, and the generator is unable to generate a sample distribution that closely matches the missing data. Secondly, the generator produces values between 0 and 1, rather than 0 or 1, which can make it challenging to interpret the actual meaning of the generated data. The main contributions of this paper are as follows:

1. We propose a new missing data filling architecture called AGAN, which is based on Generative Adversarial Networks. AGAN does not use prompt matrix but directly uses a similar discriminator network for filling, which effectively improves the filling accuracy.
2. We introduce the attribute matrix and directly convert it into real-values, two-values and nominal values. This effectively addresses the issue of the unclear interpretation of the generated data.
3. Our experimental results demonstrate that the proposed AGAN algorithm outperforms other mainstream filling algorithms in terms of lower error rates and higher downstream prediction accuracy.

### Dataset

This study utilizes two real heart disease datasets [20]. The table below provides an overview of the sample and variable characteristics of these two datasets:

The number of samples, attributes, real-values, two-values, and nominal values in these datasets differ. Conducting experiments using these data sets can reflect the performance differences of the missing data filling model in different dimensional data sets. In the case of the real heart disease data sets, it is necessary to extract the corresponding features and labels. For instance, in the Kaggle Heart dataset, heart disease is represented by “Yes”, and no disease is represented by “No”, so the corresponding “Yes” should be converted to 1, and “No” should be converted to 0. The data distribution of the two data sets is illustrated in Fig. 1.

**Fig 1.**
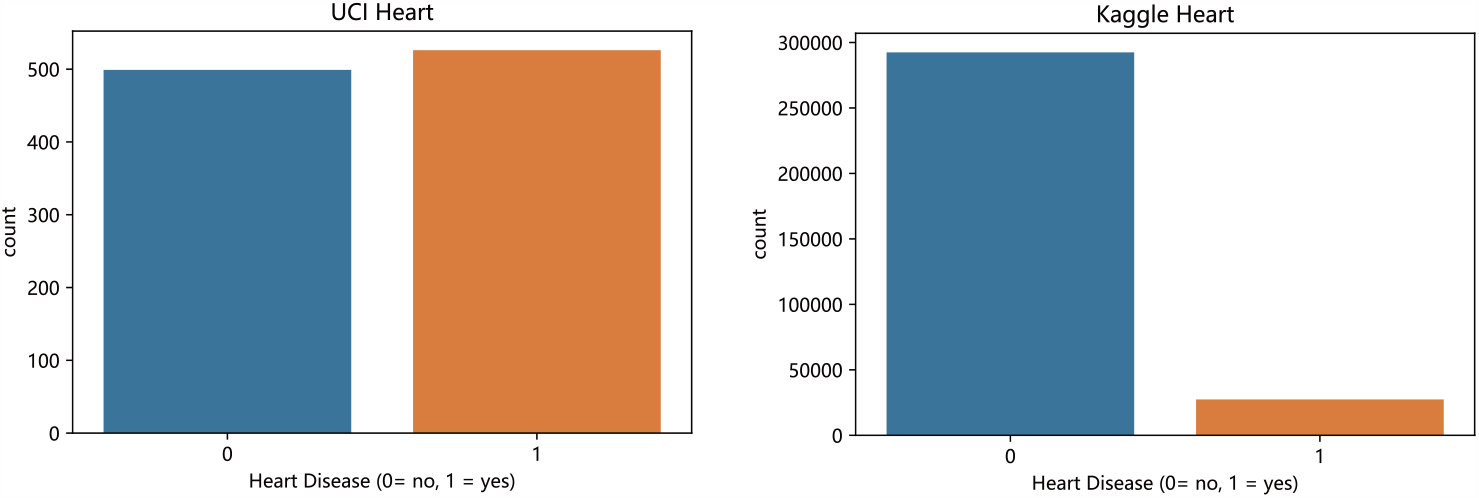
Data distribution of two real heart disease datasets

Since both datasets have a high number of features, we can analyze the correlation coefficient matrix of the two datasets to gain insight into their relationships, as shown in Fig. 2. The graph provides a visual representation of the correlation between variables, with colors indicating the strength of the relationship. Darker colors indicate weaker correlations.

**Fig 2.**
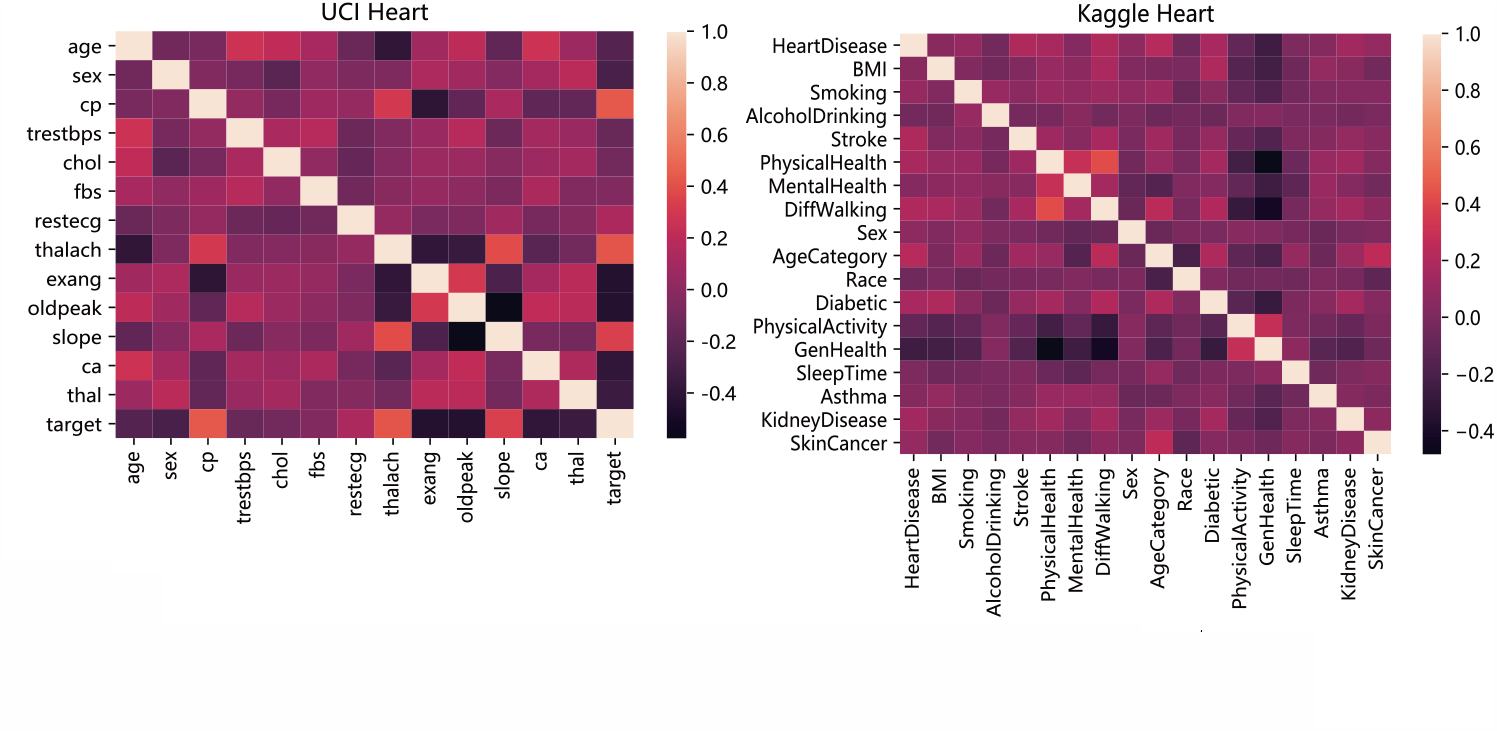
Correlation coefficient matrix of two heart disease datasets.

### UCI Heart Dataset

The UCI Heart Disease Databases was compiled by Andras Janosi, William Steinbrunn, Matthias Pfisterer, and Robert Detrano. The dataset consists of 1025 samples, each of which has 14 attribute information.

### Kaggle Heart Dataset

The Kaggle Heart dataset is sourced from the United States Centers for Disease Control and Prevention as a crucial part of the Behavioral Risk Factor Monitoring System (BRFSS), which conducts annual telephone surveys to gather health-related data of American residents. The BRFSS system undertakes over 400,000 adult interviews annually, making it the most comprehensive and continuous health survey system worldwide. This dataset includes 319,795 samples, each consisting of 18 attribute information, such as HeartDisease, BMI, Smoking, AlcoholDrinking, Stroke, PhysicalHealth, MentalHealth, DiffWalking, Sex, AgeCategory, Race, Diabetic, PhysicalActivity, GenHealth, SleepTime, Asthma, KidneyDisease, and SkinCacer.

### Methodology

#### Model Overview

This paper proposes a new missing data filling architecture AGAN based on generative adversarial networks to address the issue of diverse feature attributes in medical data, specifically in heart disease datasets [21]. The proposed method generates a feature attribute matrix for a given dataset *X* = (*X*_1_, *X*_2_, …, *X*_*n*_)^*T*^ consisting of n data samples, each with d-dimensional attribute values *X*_*i*_ = {*x*_*i*1_, *x*_*i*2_, …, *x*_*id*_} For instance, missing data on a 4*5 scale as shown in Fig. 3 can be represented as *X* = (*X*_1_, *X*_2_, *X*_3_, *X*_4_, *X*_5_)^*T*^ where “*x*_*ij*_” represents observable data, and “/” represents missing data [22].

**Fig 3.**
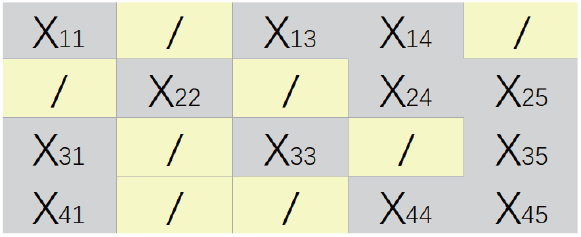
Original data example

Because the original data matrix X is incomplete, it is necessary to have another corresponding matrix to determine the information of the missing data [23]. The missing marker matrix *M* = {*m* | *m*_*ij*_ ∈ {0, 1}} is a binary matrix used to identify the location of the missing data, as shown in Fig. 4. The elements in matrix M are generated by the following formula:

**Fig 4.**
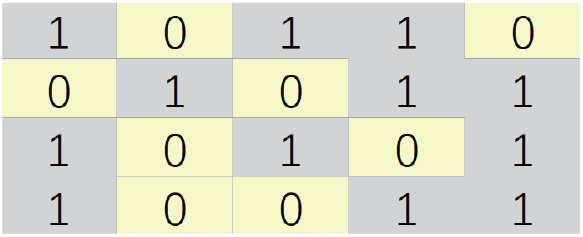
Mask data example

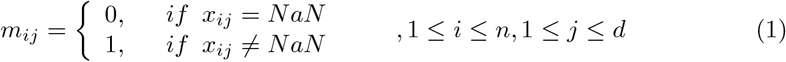

A value of 0 for *m*_*ij*_ ndicates that the jth attribute value of the ith sample is missing.

Feature attributes can be categorized into real-value type, two-value type, and nominal value type, where most of them are numerical data, while a few attributes, such as gender and whether or not, are Boolean values (0 or 1) and some are semantic discrete values such as pain type, rising and falling levels, etc. Hence, the real-value type, two-value type, and nominal value type can be represented by 0, 1, and 2, respectively [24]. Specifically, the attribute matrix is presented in Fig. 6, where matrix *A* = *{a*|*a*_*i*_ *∈ {*0, 1, 2*}}* is used to identify the attribute information of the data. The first, fourth, and fifth columns correspond to real value type, the second column corresponds to binary type, and the third column corresponds to nominal type [25].

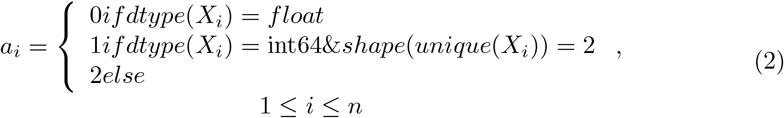

The addition of the attribute matrix provides additional information, which can constrain sample generation to a range closer to real samples and fill in more realistic data.

The overall model architecture of AGAN is shown in Fig. 5.

**Fig 5.**
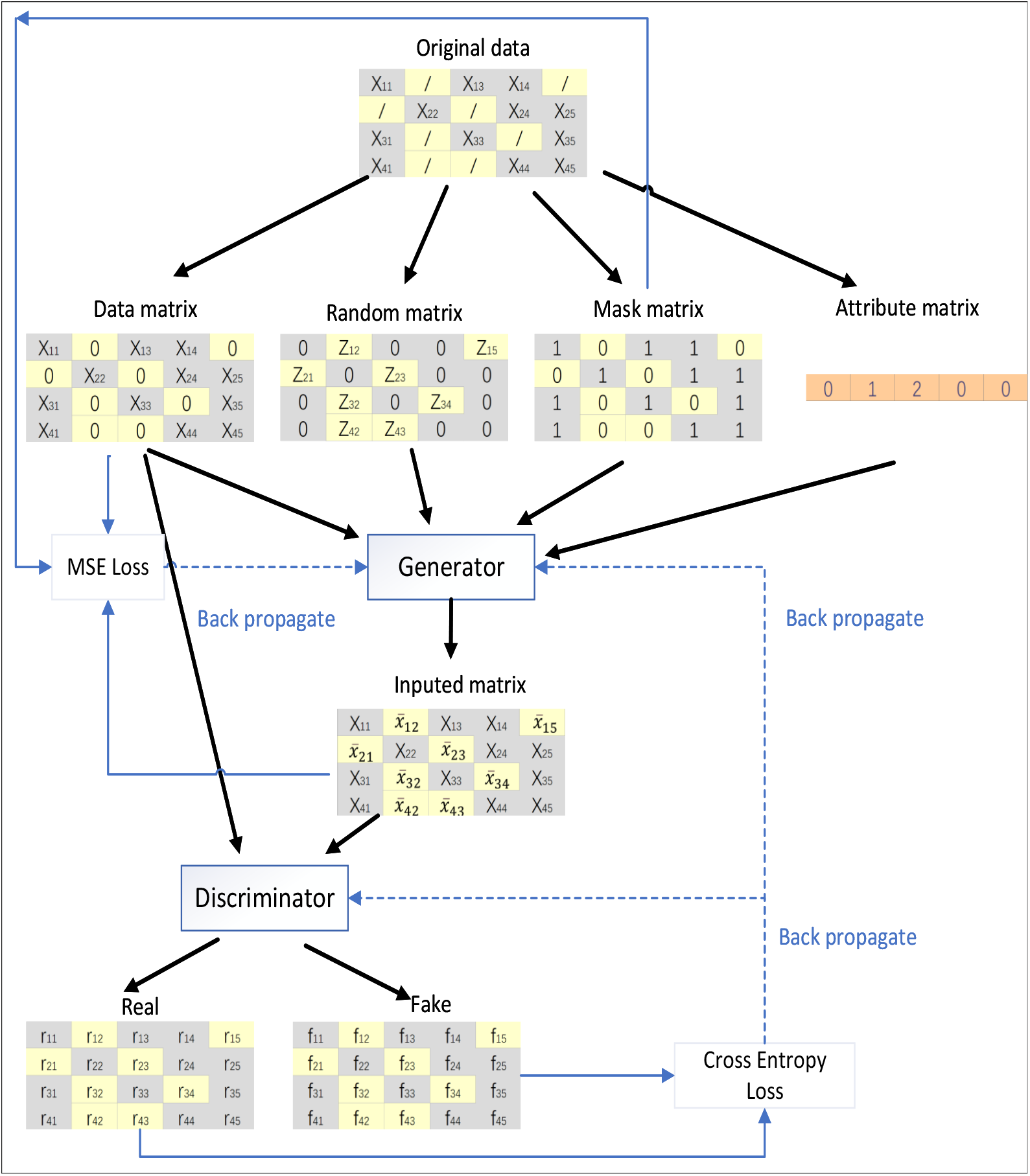
Architecture of AGAN algorithm model

**Fig 6.**
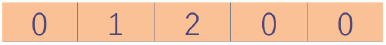
Attribute matrix example

#### Generator

The role of the generation network (represented as G) in the generative adversarial network model is to learn to fit the data distribution and generate new data. In the context of missing data filling, generating data at the missing locations is a key task. Therefore, the function of the generation network in the data filling task is to generate data. The network is generated by first fitting the distribution of real data and, in this process, learning the mapping relationship between random noise data and real data. Within the generation network, the input data includes the data matrix X, random noise data Z, marking matrix M, and attribute matrix A. The generated data is expressed in matrix 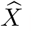. The complete data, expressed as matrix 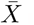, is obtained by combining the generated data obtained through the generation network with the original data that is not missing. Z is a d-dimensional vector, represented as *Z* = (*Z*_1_, *Z*_2_, …, *Z*_*d*_), where each element of 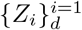 has a non-missing value in 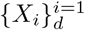, and the missing values in 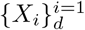 are replaced by random noise. The attribute matrix *N* = (*N*_1_, *N*_2_, …, *N*_*d*_) marks the attribute information of each column in the dataset. The function that extracts random values from a continuous uniform distribution (i.e. [-0.01, +0.01]) outputs:

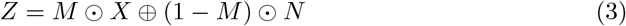

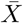 is the fill value generated by the generator, and its formula is as follows:

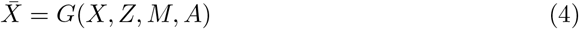

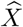 represents the final complete data. In cases where there is a missing value, the corresponding data in X is used. Conversely, where there are no missing values, the data in 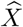 is used. Attribute matrix A is used to mark the relevant attributes. If the data is binary or nominal, it will be converted to the nearest integer value. Otherwise, the data will remain unchanged. The overall formula is as follows:

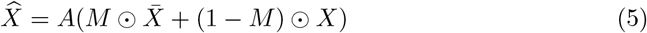

#### Discriminator

In generative adversarial networks, the discriminator network (represented as D) engages in an antagonistic game with the generator network [26]. Its main function is to determine whether the input is from real data or data generated by the generator network, and to score the input data and provide feedback to the generator network. This process is crucial in achieving the most desirable convergence state through antagonistic training, and embodies the idea of a zero-sum game.

In the missing data filling task, the function of the network is evaluated based on the data generated by the generator network. The input data is distinguished as either derived from real data or generated data from the generator network, and the differentiation result is fed back to the generator network. Consequently, the generator network adjusts its performance based on the feedback provided by the discriminator network, thereby generating data that more closely resembles real data [27].

#### Data preprocessing and model training

The first step in the missing data filling task is data preprocessing [28]. The objects of this task may include real-values, two-values, and nominal values, which cannot be directly used as input data for the filling model. Additionally, the NaN marks in incomplete data cannot be calculated. Therefore, data preprocessing is necessary for data filling. For binary and nominal attribute values, one-hot encoding is used for conversion, and NaN-marked data is replaced by the mean value of the column.

However, this method may cause missing values to be indistinguishable from real values. Therefore, another function of data preprocessing is to generate the missing marker matrix M from incomplete missing data for model training.

To prevent possible problems of gradient disappearance and gradient explosion in model training, and to enable the loss function in training to converge quickly, data normalization is performed [29]. The normalization method used in this paper is maximum and minimum normalization, which does not affect the properties of categories 0 and 1. The normalization formula is as follows:

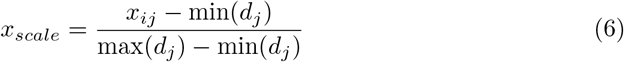

Where max(*d*_*j*_) and min(*d*_*j*_) represent the maximum and minimum values of the j-th dimension attribute, and the normalized data value *x*_*scale*_ is within the range of [0,1].

In order to solve the common problems of gradient disappearance and gradient descent in the networks, this paper introduces the variable fraction gradient descent method. There are three definitions of fractional derivatives: Riemann-Liouville, Caputo, and Grunwald-Letnikov fractional derivatives. Caputo fractional derivative [**?**], which is very suitable for engineering calculation, is used in this paper, and its definition is as follows:

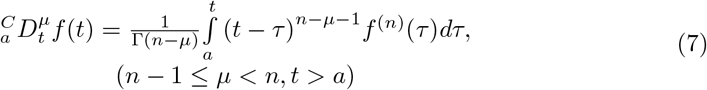

Compared to traditional methods, the iteration of the variable fraction gradient descent method can locate a better global minimum more efficiently. This translates to faster training speed and improved training accuracy compared to the traditional neural network training approach. The generator G and discriminator D typically utilize the ReLU or sigmoid function as the network activation function. However, with the ReLU function, model collapse may occur during network parameter updates, which results in neuron disappearance and the gradient flow being disrupted when the descending gradient is large. To address these issues, this section introduces a new function to modify the linear element, utilizing variable fractional order, fractional gradient descent, and gradient ascent methods in place of the traditional gradient descent or ascent methods. An alternative activation function, the Swish activation function, proposed by Google, offers improved effectiveness over ReLU [30]. Swish boasts the advantages of a lower bound, monotony, and differentiability throughout.

Moreover, Swish introduces adaptive parameters that adjust the negative slope based on specific conditions, thus enhancing activation efficiency. It is defined as:

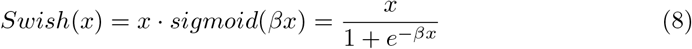

Where *β* represents the adaptive parameter. As the positive half-axis of the Swish function is linear, the curves of the first and second derivatives are smooth, making gradient calculation straightforward. Consequently, the operational cost of the neural network is reduced, and its operational efficiency is improved.

During network training, the generator and discriminator are cross-trained, with the discriminator trained k times every time the generator is trained in a confrontation [31]. To train the generator, the discriminator parameters are fixed and connected in series to the generator. The prediction loss of the generator is then obtained from the difference between the filled data generated by the generator and the real data. The output probability value of the discriminator is used to calculate the confrontation loss of the generator, and the gradient information of the generator network parameters is computed by the loss function. The parameters are then updated along the negative gradient direction. When training the discriminator, the parameters of the generator are fixed, and false filling data are generated. The discriminator judges and outputs the probability that the current input data originates from the real sample set. The overall network training algorithm is presented in Table 2.

**Table 1.** Overview of attribute information of real heart disease datasets.

| Dataset | sample size | features | real-values | two-values | nominal values |
| --- | --- | --- | --- | --- | --- |
| UCI Heart | 1025 | 14 | 6 | 4 | 4 |
| Kaggle Heart | 319795 | 18 | 5 | 11 | 2 |

**Table 2.**
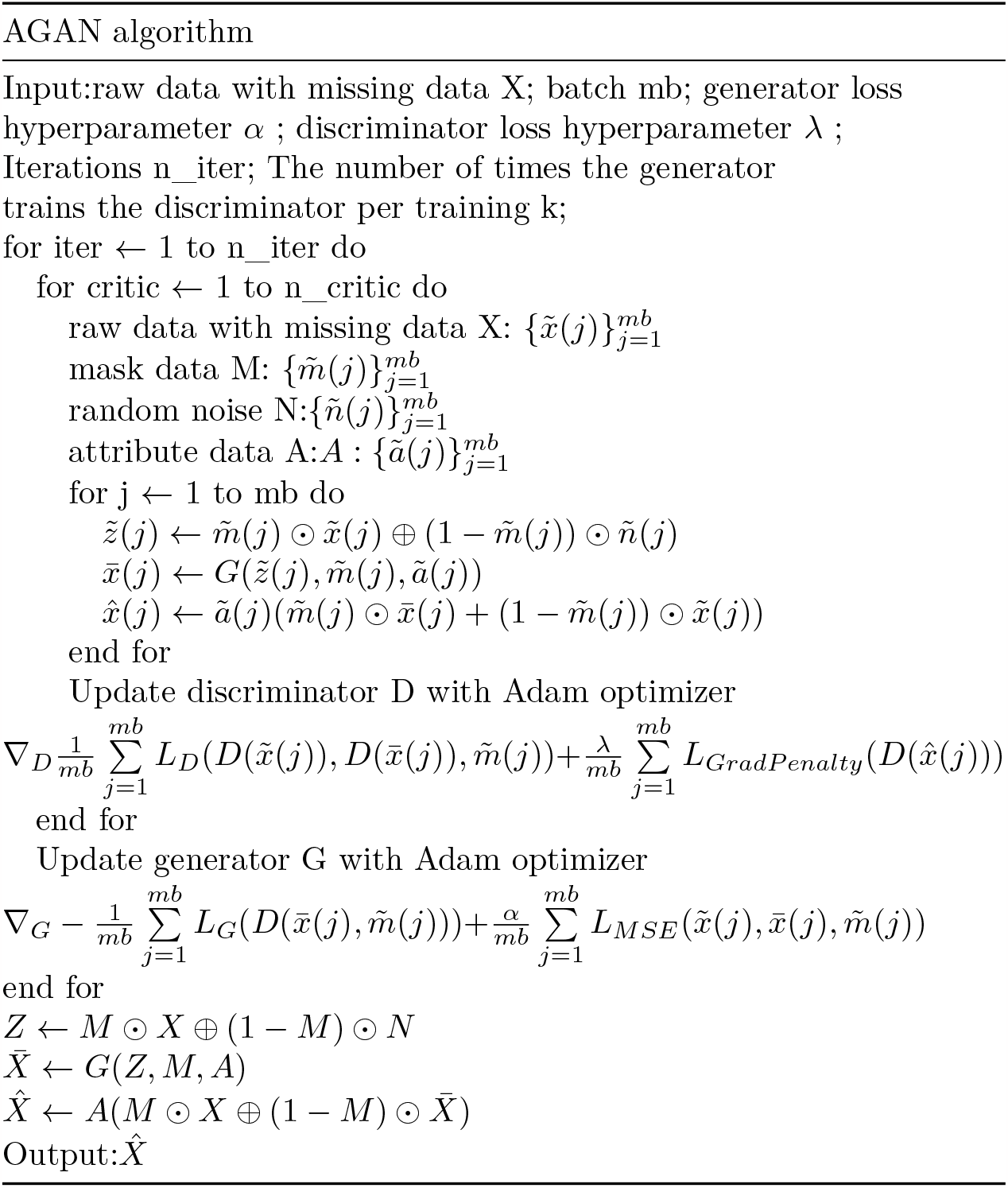
Training algorithm flow.

### Experiments & discussion

#### Experimental environment

The hardware and software environment utilized in this experiment is shown in Table 3.

**Table 3.** Experimental hardware and software environment.

| environment | value |
| --- | --- |
| CPU | Intel(R) Core(TM) i9-10920X<br>CPU@3.50GHz |
| harddisk | 1T SSD |
| RAM | 128GB |
| OS | Windows 10 |
| program language | Python 3.7 |
| deep learning framework | Pytorch 1.10 |
| GPU | NVIDIA GeForce RTX3090 |

#### Performance metrics

1. Root mean square error (RMSE) The root mean square error (RMSE) is the evaluation index used in this paper to assess the experimental results. RMSE measures the difference between data and is utilized in this experiment to calculate the error between the real and filled data. The calculation formula for RMSE is as follows:

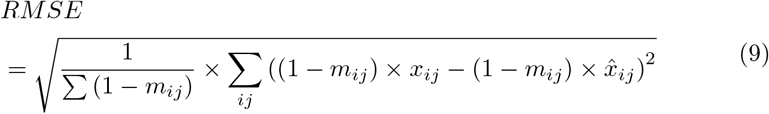

The RMSE utilized to evaluate the filling effect in this experiment only calculates the error between the missing value and its corresponding filling value, and does not factor in the observed data in the data sample during the calculation. This approach helps avoid the impact of different missing rates on the filling evaluation.
2. Proportion of falsely classified (PFC) The second evaluation index used in this paper is PFC, which is the performance index for category variables. A smaller value and a smaller error indicate a better filling effect. The calculation formula for PFC is as follows:

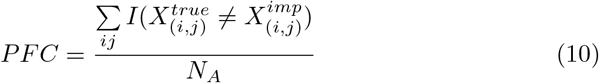

In the above equation, 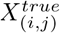 represents the value of the j-th variable of the i-th sample in the complete dataset, 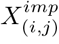 represents the value of the j-th variable of the i-th sample in the filled dataset, and *N*_*A*_is the number of missing discrete variables.
3. Area Under Curve and receiver operating characteristic curve This paper utilizes the AUC (Area Under Curve) index to reflect the indirect comparison between the two classification models used in the experiment. AUC represents the area under the subject’s operating curve. The AUC index can assess the quality of the classifier even when the sample label distribution is uneven. The calculation of AUC is dependent on the number of positive true positive rates (TPR) and false positive rates (FPR). The calculation formula for FPR and TPR is shown below:

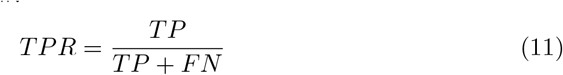

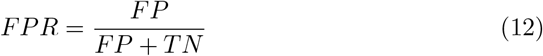

By plotting FPR as the ordinate and TPR as the abscissa, the receiver operating characteristic curve can be generated. AUC is the area under this curve, i.e.:

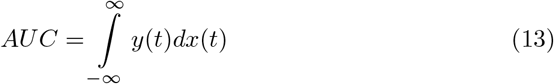

Where x and y are the horizontal and vertical coordinates of the curve in the figure. AUC represents the probability of a classifier ranking a randomly chosen positive sample higher than a randomly chosen negative sample. In a binary classification model, a higher AUC indicates that the classifier has better discrimination power between positive and negative samples. The AUC index is not affected by the distribution proportion of positive and negative samples, making it a useful metric in situations where the sample size or class distribution is imbalanced. Compared to metrics such as recall and accuracy, AUC can more accurately assess the quality of a classification model in such situations.

#### Experimental results and discussion

In order to evaluate the performance of the proposed missing data filling method, a complete dataset without any missing values is used. The missing data is artificially generated by randomly selecting a subset of values from the original dataset and replacing them with the missing marker “NaN” to obtain the incomplete dataset to be filled. The amount of missing data in the incomplete dataset is determined by the missing rate, which is varied between 10% and 60% in this experiment. The complete random missing mechanism is used to ensure that the missing values are randomly distributed and avoid any bias towards specific attributes or samples.

To verify the filling effect of the proposed model, several rounds of tests were conducted on two types of heart disease datasets under the same experimental conditions. Comparative experiments and analyses were carried out using commonly used filling algorithms and models. The experiments included filling tests with different filling methods at various missing rates, and the resulting data were evaluated using RMSE, PFC, and AUC indicators. The differences in experimental results were briefly analyzed. Fig. 7 shows the loss of the generator and discriminator in the experiment. In these experiments, various methods are employed to fill the missing heart disease data. The filling methods include zero imputation, missing forest filling [32], multiple imputation by chained equations (MICE), K nearest neighbor(KNN), GAIN model, and AGAN, which is the filling algorithm proposed in this paper. AGAN and GAIN are generative machine learning models, while MissForest, MICE, and KNN are discriminative machine learning models [33]. Zero imputation is used as a reference for comparison.

**Fig 7.**
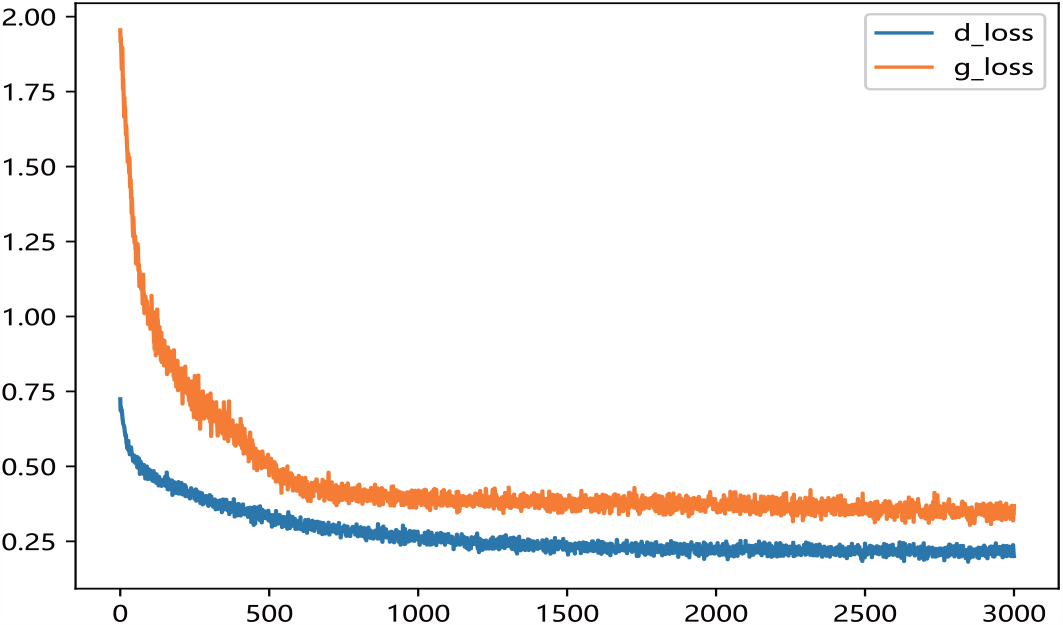
Loss of generator and discriminator during training

1. RMSE of missing data filling algorithms on different datasets This experiment compares the filling effects of six different methods on two datasets, and Table 4 shows the root mean square error of filling for each method. The experiment utilizes data with a missing rate of 30%. Table 4 demonstrates the notable influence of data dimension and features on the RMSE filled with missing data. Both experimental datasets possess distinct data dimensions and features. In the case of UCI Heart, a dataset with low data dimensions and few attributes, AGAN showcases the best filling effect, with GAIN exhibiting a minor disparity from the discriminant machine learning model. For instance, the filling effect of MissForest slightly surpasses that of GAIN. Conversely, on the dataset Kaggle Heart, which has higher data dimensions and more attributes, AGAN exhibits the smallest error, with an RMSE value of 0.1095, approximately 26% lower than that of the GAIN model and about 30% lower than that of MissForest, which displays the best effect in the discriminant model.

**Table 4.** RMSE value of missing data filling algorithms on different datasets (mean *±* std)
2. RMSE of missing data filling algorithms under different missing rates This experiment conducted a comparison of the filling effect of six algorithms on the UCI Heart dataset. The root mean squared error RMSE of the filling is presented in Table 5. The experiment employed data with missing rates ranging from 10% to 60%. For ease of display, a visualization diagram depicting the table is presented in Fig. 8.

| algorithm | UCI Heart | Kaggle Heart |
| --- | --- | --- |
| zero imputation | 0.7481±.0063 | 0.5862±.0049 |
| KNN | 0.2419±.0050 | 0.2033±.0034 |
| MissForest | 0.1637±.0057 | 0.1580±.0020 |
| MICE | 0.2943±.0021 | 0.2649±.0032 |
| GAIN | 0.1752±.0036 | 0.1481±.0012 |
| AGAN | 0.1284±.0011 | 0.1095±.0028 |

**Table 5.** RMSE value of missing data filling algorithms on different datasets (mean *±* std)

| algorithm | 10% | 20% | 40% | 60% |
| --- | --- | --- | --- | --- |
| zero imputation | 0.7474±.00032 | 0.7472±.0029 | 0.7527±.0048 | 0.7657±.0034 |
| KNN | 0.1263±.0012 | 0.1933±.0003 | 0.3101±.0080 | 0.5340±.0049 |
| MissForest | 0.0768±.0036 | 0.1329±.0035 | 0.2676±.0033 | 0.4205±.0061 |
| MICE | 0.2540±.0019 | 0.3105±.0011 | 0.4367±.0045 | 0.6076±.0051 |
| GAIN | 0.0727±.0030 | 0.1491±.0005 | 0.2569±.0078 | 0.3181±.0063 |
| AGAN | 0.0814±.0021 | 0.1269±.0014 | 0.1462±.0026 | 0.2371±.0057 |

**Fig 8.**
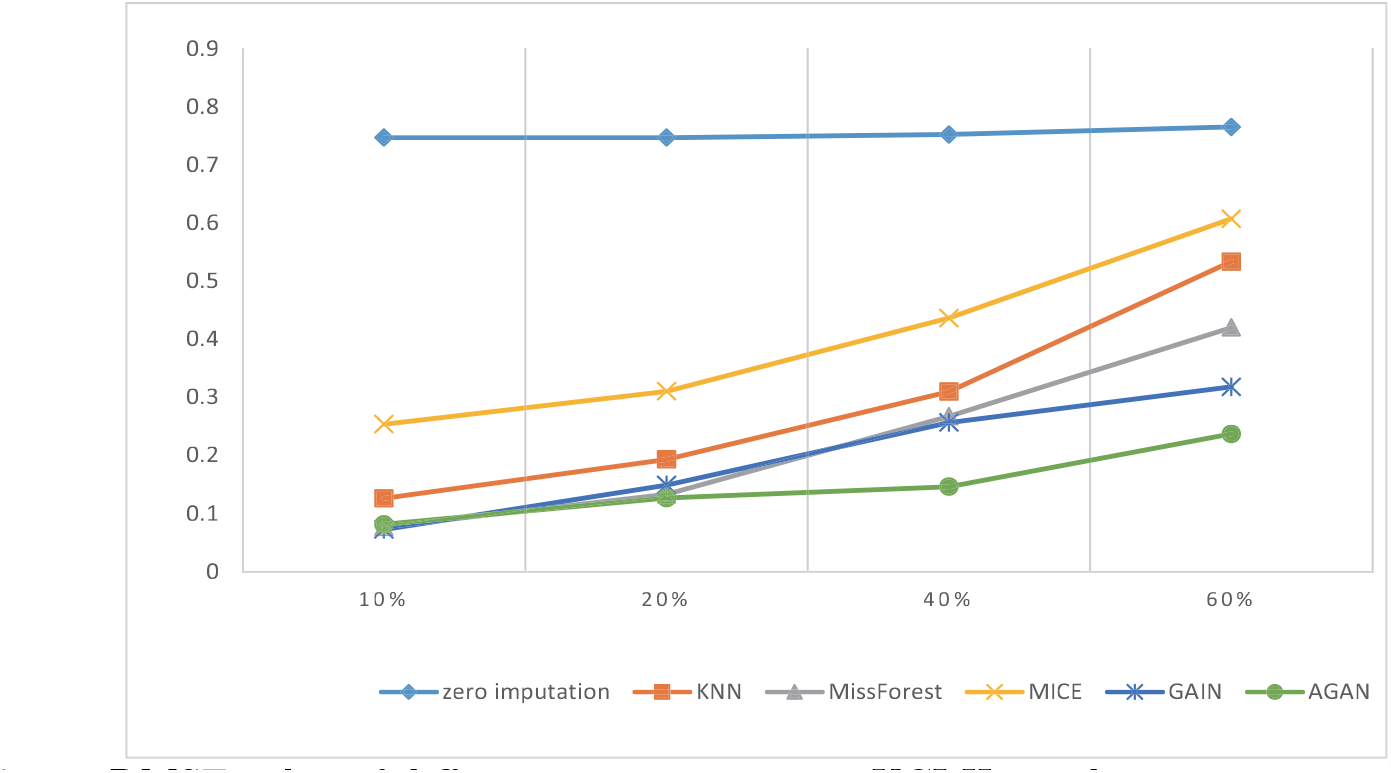
RMSE value of different missing rates on UCI Heart dataset The visualization diagram in Fig. 8 demonstrates that the filled RMSE error gradually increases with the increase of the missing rate. At a 10% deletion rate, AGAN exhibits an RMSE value of only 0.0814. MissForest, GAIN, and AGAN show the smallest filling error at low missing rates. As the missing rate exceeds 40%, the filling effect error of the discriminant algorithm increases significantly, while the algorithm based on the generation of confrontation network shows a slower increase in RMSE value. Compared to GAIN, the AGAN proposed in this study achieves the best filling effect and the smallest error.
3. RMSE of missing data filling algorithms in different sizes This experiment compares the filling effect of six algorithms on Kaggle Heart datasets of varying sizes, ranging from 5000 to 30000. Table 6 shows the filled root mean square error and standard deviation. The missing rate of the data used in this experiment is 30%. When controlling for the number of features and the missing proportion of control variables, specifically a number of 17 variables and a 30% missing proportion, the result spresented in the table are obtained by calculating four datasets of varying sizes. The table reveals that the RMSE of the four missing values filled by AGAN is the smallest, indicating the best filling effect. GAIN and MissForest follow AGAN in their filling effect, while the MICE algorithm performs the worst under these conditions. When looking at the RMSE value, GAIN’s effect is similar to that of MissForest, while the effect of AGAN is approximately 24% higher than that of GAIN, and approximately 83% higher than that of the reference algorithm.

**Table 6.** RMSE value filled by different algorithms (mean *±* std)
4. PFC of missing data filling algorithms on different datasets This experiment compares the PFC of six methods on UCI Heart and Kaggle Heart datasets. The PFC is shown in Table 7, and the missing rate of the data used in this experiment is 30%. The PFC indicator measures the proportion of missing category variables that are not accurately represented in the reconstructed data. For instance, in the table, the first data point (0.2577) indicates that 25.77% of the missing category variables were filled with an incorrect category using the zero-imputation method. A more precise PFC indicator correlates with a better quality of data reconstruction. It is worth noting that the PFC index for category variables demonstrates that the filling ability of the six imputation algorithms, including zero imputation, MissForest, KNN, MICE, GAIN, and AGAN for missing data, increases in that order. This observation aligns with the results obtained from the evaluation of the RMSE.

| algorithm | 5000*17 | 10000*17 | 20000*17 | 30000*17 |
| --- | --- | --- | --- | --- |
| zero imputation | 0.7024±.0039 | 0.6243±.0059 | 0.6747±.0062 | 0.6065±.0048 |
| KNN | 0.2389±.0012 | 0.2032±.0003 | 0.2139±.0080 | 0.2165±.0037 |
| MissForest | 0.1562±.0028 | 0.1781±.0052 | 0.1591±.0037 | 0.1619±.0021 |
| MICE | 0.2494±.0010 | 0.2817±.0031 | 0.2588±.0037 | 0.2576±.0029 |
| GAIN | 0.1697±.0021 | 0.1494±.0025 | 0.1737±.0090 | 0.1773±.0011 |
| AGAN | 0.1250±.0027 | 0.1241±.0034 | 0.1585±.0013 | 0.1363±.0014 |

**Table 7.** PFC value of algorithms on two datasets.
5. AUC of downstream classification tasks with 20% missing rate Table 8 presents the AUC values for the downstream classification task with a missing rate of 20%. To enhance the visual representation of the results, the data in the table is presented in the form of a histogram, illustrated in Figure 8. Figure 9 displays the ROC curve of the UCI Heart dataset with missing rate of 20% Table 9 presents the AUC values of the complete dataset filled using different algorithms with various classification algorithms for downstream classification tasks. The classification effect improves with a larger AUC value. The ROC curve in Figure 9 illustrates that the classification effect of the filling algorithm based on generative adversarial networks surpasses that based on discriminant. Furthermore, the proposed AGAN model in this study outperforms previous algorithms.

| algorithm | UCI Heart | Kaggle Heart |
| --- | --- | --- |
| zero imputation | 0.2577 | 0.2682 |
| KNN | 0.1737 | 0.1773 |
| MissForest | 0.1312 | 0.1427 |
| MICE | 0.1854 | 0.2055 |
| GAIN | 0.1230 | 0.1415 |
| AGAN | 0.0912 | 0.1013 |

**Table 8.** AUC indicators of classified tasks.

| algorithm | UCI Heart | Kaggle Heart |
| --- | --- | --- |
| zero imputation | 0.7445 | 0.6598 |
| KNN | 0.8042 | 0.7542 |
| MissForest | 0.8221 | 0.7626 |
| MICE | 0.7279 | 0.6922 |
| GAIN | 0.8359 | 0.7819 |
| AGAN | 0.8706 | 0.8085 |

**Fig 9.**
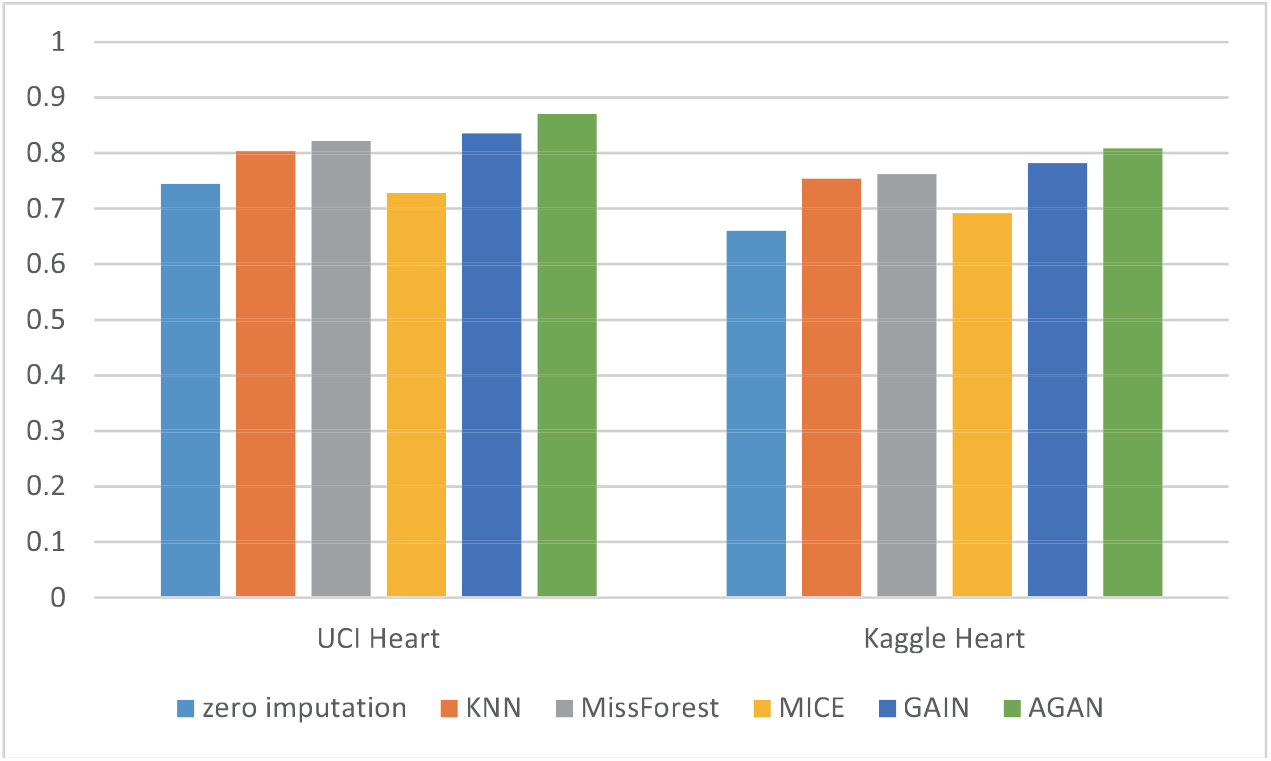
Histogram of classification the AUC indicators for classification after imputing the heart disease dataset

**Fig 10.**
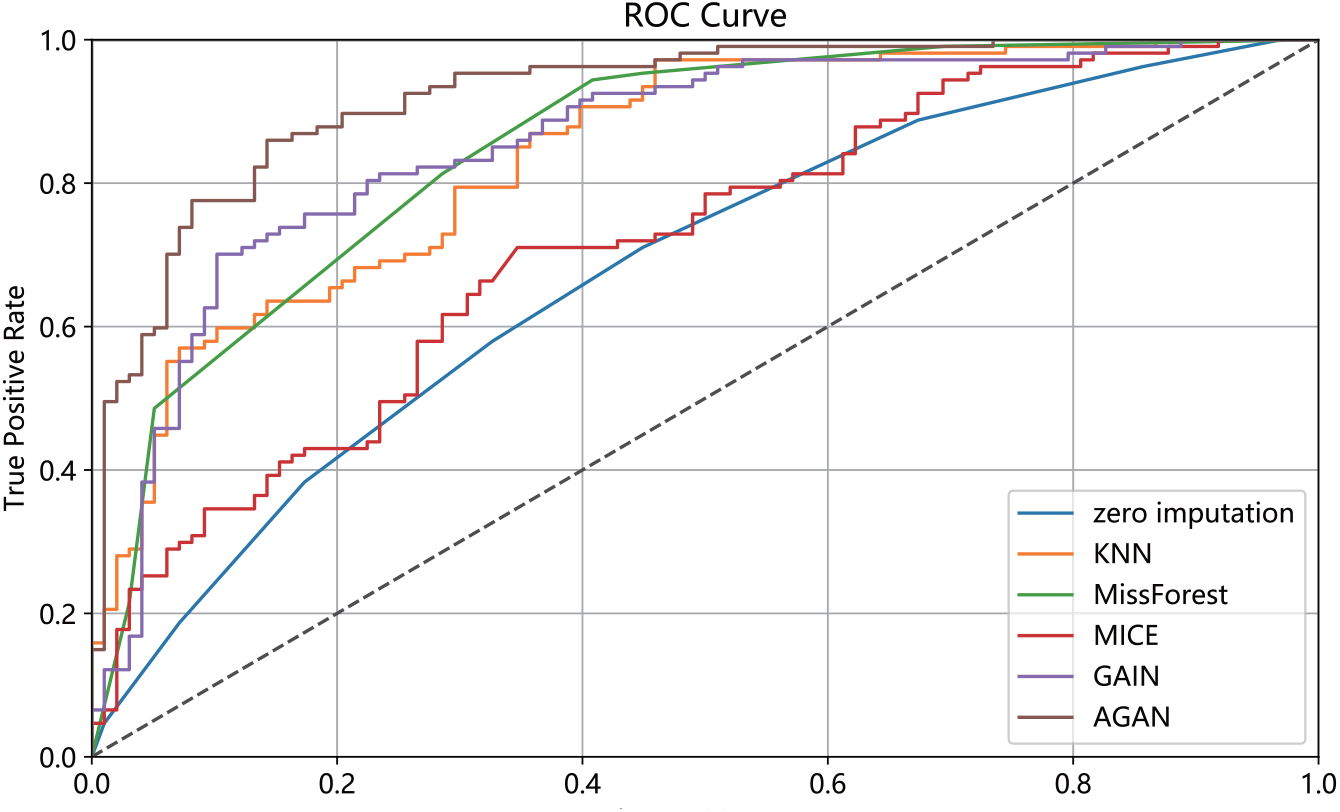
Figure 9 ROC curves of six algorithms

## Conclusion

In many domains, including medicine, missing data is a common issue in datasets. In this paper, we propose a algorithm for filling missing data in cardiac disease datasets using generative adversarial networks. Our approach involves introducing an attribute matrix to aid in filling in missing variables, and using the Swish activation function to address gradient disappearance and explosion during training. We conducted six experiments to evaluate the proposed model and analyzed the results. Our findings show that our missing data filling algorithm based on generative adversarial networks outperforms the missing data filling algorithm based on discriminant, as evidenced by significantly better RMSE and AUC. The error rate of our algorithm is over 10% lower than that of the experimental comparison algorithm. Additionally, our approach achieves the lowest RMSE and highest AUC in the experimental comparison algorithm. We also observe that as the missing rate increases, the growth of RMSE is relatively slow. Future research can focus on improving network sparsity algorithm s and optimizing neural network structures based on attribute correlations to enhance interpretability and gap filling. Other potential areas of research include incorporating convolution, BatchNormal, and other technical means to improve network efficiency.

## Acknowledgments

This work is supported by the Jilin Scientific and Technological Development Program 20210402078GH.

